# Transfer RNA genes experience exceptionally elevated mutation rates

**DOI:** 10.1101/229906

**Authors:** Bryan P. Thornlow, Josh Hough, Jacquelyn M. Roger, Henry Gong, Todd M. Lowe, Russell B. Corbett-Detig

**Author notes:** H.G. performed initial analyses. B.P.T. performed research and analyzed data, with assistance from J.M.R. J.H. performed the DFE analyses. B.P.T., T.M.L. and R.B.C.-D. conceived and designed research. B.P.T., T.M.L. and R.B.C.-D. wrote the paper. J.H., J.M.R., H.G., T.M.L. and R.B.C.-D. edited the paper.

## Abstract

Transfer RNAs (tRNAs) are a central component for the biological synthesis of proteins, and they are among the most highly conserved and frequently transcribed genes in all living things. Despite their clear significance for fundamental cellular processes, the forces governing tRNA evolution are poorly understood. We present evidence that transcription-associated mutagenesis and strong purifying selection are key determinants of patterns of sequence variation within and surrounding tRNA genes in humans and diverse model organisms. Remarkably, the mutation rate at broadly expressed cytosolic tRNA loci is likely between seven and ten times greater than the nuclear genome average. Furthermore, evolutionary analyses provide strong evidence that tRNA genes, but not their flanking sequences, experience strong purifying selection, acting against this elevated mutation rate. We also find a strong correlation between tRNA expression levels and the mutation rates in their immediate flanking regions, suggesting a simple new method for estimating individual tRNA gene activity. Collectively, this study illuminates the extreme competing forces in tRNA gene evolution, and implies that mutations at tRNA loci contribute disproportionately to mutational load and have unexplored fitness consequences in human populations.

**Significance Statement:** While transcription-associated mutagenesis (TAM) has been demonstrated for protein coding genes, its implications in shaping genome structure at transfer RNA (tRNA) loci in metazoans have not been fully appreciated. We show that cytosolic tRNAs are a striking example of TAM because of their variable rates of transcription, well-defined boundaries and internal promoter sequences. tRNA loci have a mutation rate approximately seven-to tenfold greater than the genome-wide average, and these mutations are consistent with signatures of TAM. These observations indicate that tRNA loci are disproportionately large contributors to mutational load in the human genome. Furthermore, the correlations between tRNA locus variation and transcription implicate that prediction of tRNA gene expression based on sequence variation data is possible.

---

Transfer RNAs (tRNAs) are essential to protein synthesis across all of life. Their primary function is in translation of the genetic code into the corresponding amino acid sequences that make up proteins. Thus, tRNA molecules are critical for virtually all cellular processes, and the genes encoding tRNA molecules have been highly conserved over evolutionary time (1, 2). Mitochondrial tRNAs have been the subject of many studies, as mutations in these genes lead to a large number of maternally inherited genetic diseases (3). However, eukaryotic genomes contain approximately 10-to 20-fold as many tRNA genes encoded in their nuclear chromosomes, which are required for cytosolic protein translation (2, 4). In spite of their importance to the cell, there has been little study of evolutionary conservation or pathogenic mutations in cytosolic tRNA genes (5, 6). tRNAs are required in exceptionally large quantities, and therefore tRNA genes may experience greater levels of transcription than even the most highly transcribed protein-coding genes (7, 8). In turn, this may lead to high levels of transcription-associated mutagenesis (TAM). As the largest, most ubiquitous RNA gene family, cytosolic tRNAs constitute an ideal gene set for studying the interplay between natural selection and elevated mutation rates.

Transcription affects the mutation rates of transcribed genes (9) through the unwinding and separation of complementary DNA strands (10). During transcription, a nascent RNA strand forms a hybrid DNA-RNA complex with a template DNA strand. While the complementary tract of non-template DNA is temporarily isolated, it is chemically reactive and thus accessible by potential mutagens (10). Transcription can lead to the formation of non-canonical DNA structures, which can hinder repair pathways and promote errors by the polymerase (11). The RNA strand can also re-anneal to the template DNA strand, prolonging isolation and increasing vulnerability to mutations (12, 13). Furthermore, if transcription and DNA replication occur concomitantly at a particular locus, collisions between RNA polymerase and the DNA replication fork may also damage DNA (9, 11, 14). In human cancer cells, increased transcription and replication induces torsional stress and collisions (11).

Several cellular agents have also been shown to cause damage in highly expressed genes (15). Among the most notable sources of mutation associated with transcription is activation-induced cytidine deaminase (AID; 16). AID accompanies RNA polymerase II and deaminates cytosine nucleotides. To resolve the resulting base-pair mismatch, the opposing guanine is converted to adenine and uracil to thymine, resulting in excess C→T mutations on the non-template strand and excess G→A mutations on the template strand (9, 17). AID is a member of the APOBEC gene family, many of which are involved in double-stranded break repair in transcription (9). Some members of the APOBEC family act strongly at short genes, suggesting increased activity at tRNA loci (18, 19). For example, APOBEC3B causes 1000-fold more DNA damage at tRNA loci than at other genomic regions in yeast (19). AID also acts on highly transcribed genes in immune B-cells, causing transition mutations and double-stranded breaks (9). Due to the strong association of the APOBEC family with transcription, relative excesses of C→T and G→A mutations are a signature of TAM (9).

To conserve mature tRNA sequence identity in the presence of an elevated mutation rate, tRNA genes should experience strong purifying selection. tRNA transcription requires sequence-specific binding of transcription factors to the internal box A and box B promoter elements (20). Once transcribed, precursor tRNAs must fold properly to undergo maturation, which can be disrupted by sequence-altering mutations. The unique structure of tRNAs dictates processing by RNases, addition of modifications, accurate recognition by aminoacyl tRNA synthetases, incorporation into the translating ribosome, and accurate positioning of the anticodon relative to mRNA codons (21, 22). Because of the need to maintain sequence specificity, DNA encoding the mature portions of tRNAs are well conserved (21). Therefore, we expect that a large proportion of mutations arising in tRNA genes will be deleterious and will quickly be purged by natural selection.

While most human tRNA genes do not have external promoters (20, 21), tRNA transcripts include leader and trailer sequences, extending roughly 2-5 nucleotides upstream and 5-15 nucleotides downstream of the annotated mature tRNA gene, based on the position of the genomically encoded poly(T) transcription termination sequence. Aside from the termination sequence, these flanking sequences appear to have limited sequence-specific functionality in most cases (23–26). Very early in maturation, all tRNA flanking sequences are removed by RNase P (22–24) and RNase Z (22, 27). Because these flanking genomic sequences are frequently unwound and therefore vulnerable to TAM, we expect that these regions will experience similar mutation rates to tRNAs. Whereas tRNA genes should experience purifying selection, the flanking regions should be neutral or under weak selection. Here we investigate the patterns of conservation, divergence and within-species variation of cytosolic tRNAs in humans and other model organisms to elucidate the forces shaping the evolution of this essential RNA gene family.

## Results & Discussion

### Flanking regions of tRNA genes are highly variable despite strong conservation of mature tRNA sequences

To estimate evolutionary conservation, we examined PhyloP, which measures conservation of each human genomic position across 100 vertebrate species (28), by position within each tRNA locus (see Methods). Positive PhyloP scores indicate strong conservation and negative scores indicate accelerated evolution. To study the effects of evolution on a shorter time-scale, we also estimated sequence divergence between human and *Macacca mulatta* at each tRNA locus. Mature tRNA sequences are highly conserved across all positions, based on both average PhyloP score (28; Fig. 1A; SI Appendix, Dataset S1) and *M. mulatta* alignment (Fig. 1B). However, the inner 5’ flanking region (20 bases upstream of the tRNA, see Methods) is roughly four times more divergent than the untranscribed reference regions. We also find increased rates of divergence in the inner 3’ flanking region, which is roughly three times more divergent than the reference regions (Fig. 1B). Both the outer 5’ flank (21-40 bases upstream of the tRNA) and outer 3’ flank (11-40 bases downstream) are also roughly 1.5 times more divergent than the reference regions. For tRNAs that contain introns (2), we find that intronic variation correlates with flanking variation (SI Appendix, Fig. S1). Furthermore, intergenic regions within clusters of active tRNAs show similar patterns in their PhyloP scores (SI Appendix, Fig. S2).

**Fig. 1.**
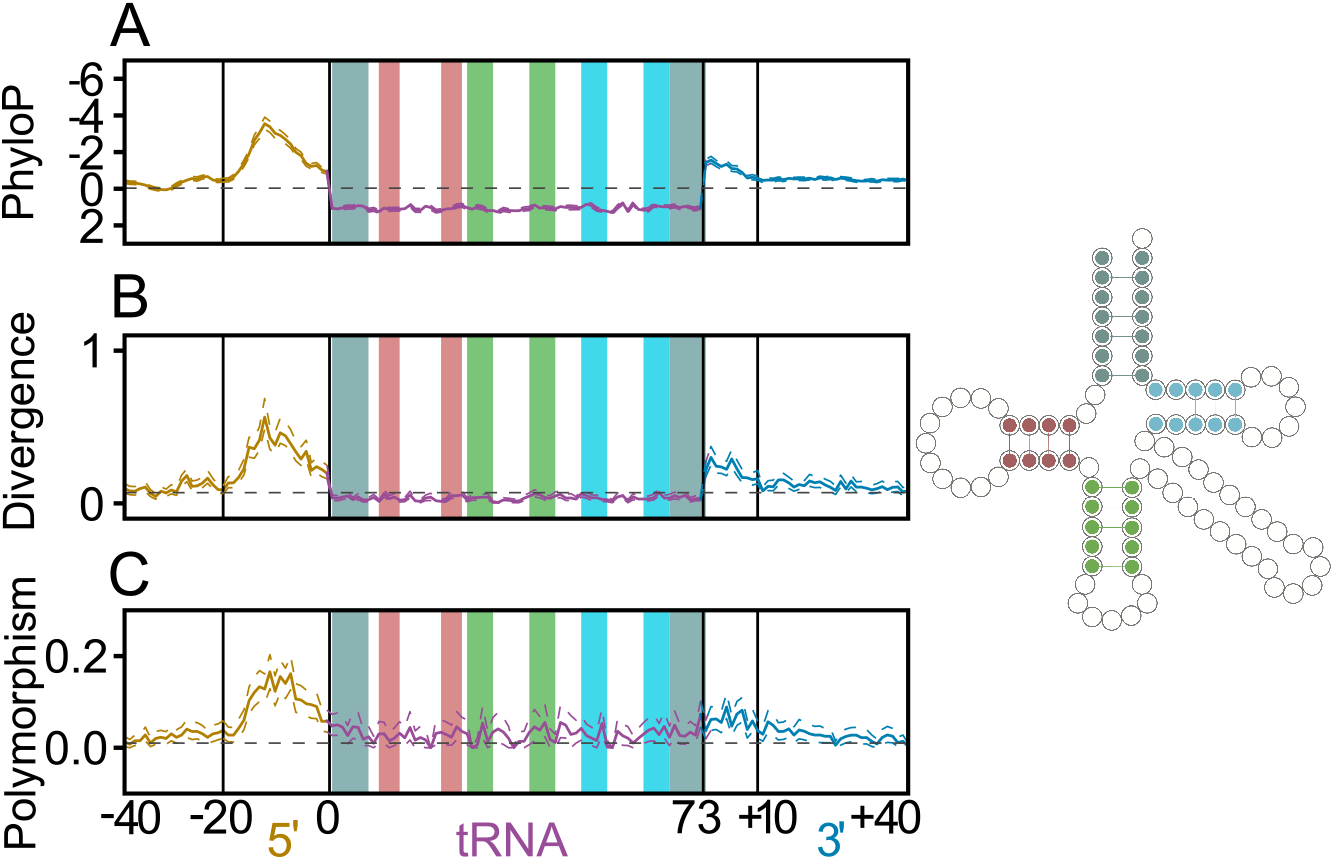
Strong pattern of variation in regions flanking human tRNA genes relative to vertebrates, upon comparison to Rhesus macaque, and within the human population. (**A**) Average PhyloP score (comparing humans to 100 vertebrate species) is plotted for each position within the tRNA and flank, across all human tRNAs. (**B**) Divergence between the human and *M. mulatta* tRNA genes and their flanking regions. (**C**) Frequency of low-frequency SNPs (minor allele frequency less than or equal to 0.05) across all human tRNAs. The acceptor stem (gray), D-stem (red), anticodon stem (green) and T-stem (blue) are highlighted within the tRNA, both in the linear plots and in the 2D structure legend to the right (2, 31). Nucleotide numbering below plot relative to mature tRNA boundaries, with inner and outer flanks demarcated by a shift in mutation rate (see Methods). Dotted lines surrounding plots depict 95% confidence intervals, calculated by non-parametric bootstrapping by tRNA loci.

We also studied population-level variation at low-frequency single nucleotide polymorphisms (SNPs; minor allele frequency < 0.05%) for each tRNA locus. Low-frequency SNPs are evolutionarily young, and less affected by selection (29). Consistent with our divergence analyses, we find that low-frequency SNPs are more common across both the tRNA gene sequence and flanking regions than in untranscribed reference regions (Fig. 1C). Although the inner flanking regions are most polymorphic, the mature tRNA sequences have about twice as many low-frequency SNPs as reference regions. Overall, our results are consistent on multiple timescales, indicating that tRNAs and flanking sequences are prone to mutation. Indeed, of the 247 sites in the genome that have the lowest possible PhyloP scores, -20 (28, 30), 14 are 10 to 15 bases upstream of the start of an active tRNA gene, indicating disproportionate enrichment (hypergeometric test, p < 1.65e-48) and that tRNA flanking regions are among the least conserved in the genome. Nonetheless, mature tRNA gene sequences are strongly conserved by purifying selection, which purges mutations.

### Transcription is correlated with variation in tRNA and flanking regions

We hypothesized that, if transcription-associated mutagenesis drives variation among tRNA loci, highly active tRNA genes would show the greatest mutation rates. Because tRNA transcript abundance measures are often not attributable to individual loci due to identical gene copies and difficulty sequencing full-length tRNAs, we estimated relative transcriptional activity based on chromatin state data from the Epigenomic Roadmap Project (33). Based on these data, we classified human tRNA genes as “active” if they are located in expressed regions in several cell lines and otherwise “inactive” (Fig. 2; see Methods). In some cases, multiple cell lines correspond to a single tissue or organ, so tissue-specific tRNAs (e.g. the brain-specific arginine tRNA in mouse (6)) are considered active.

**Fig. 2.**
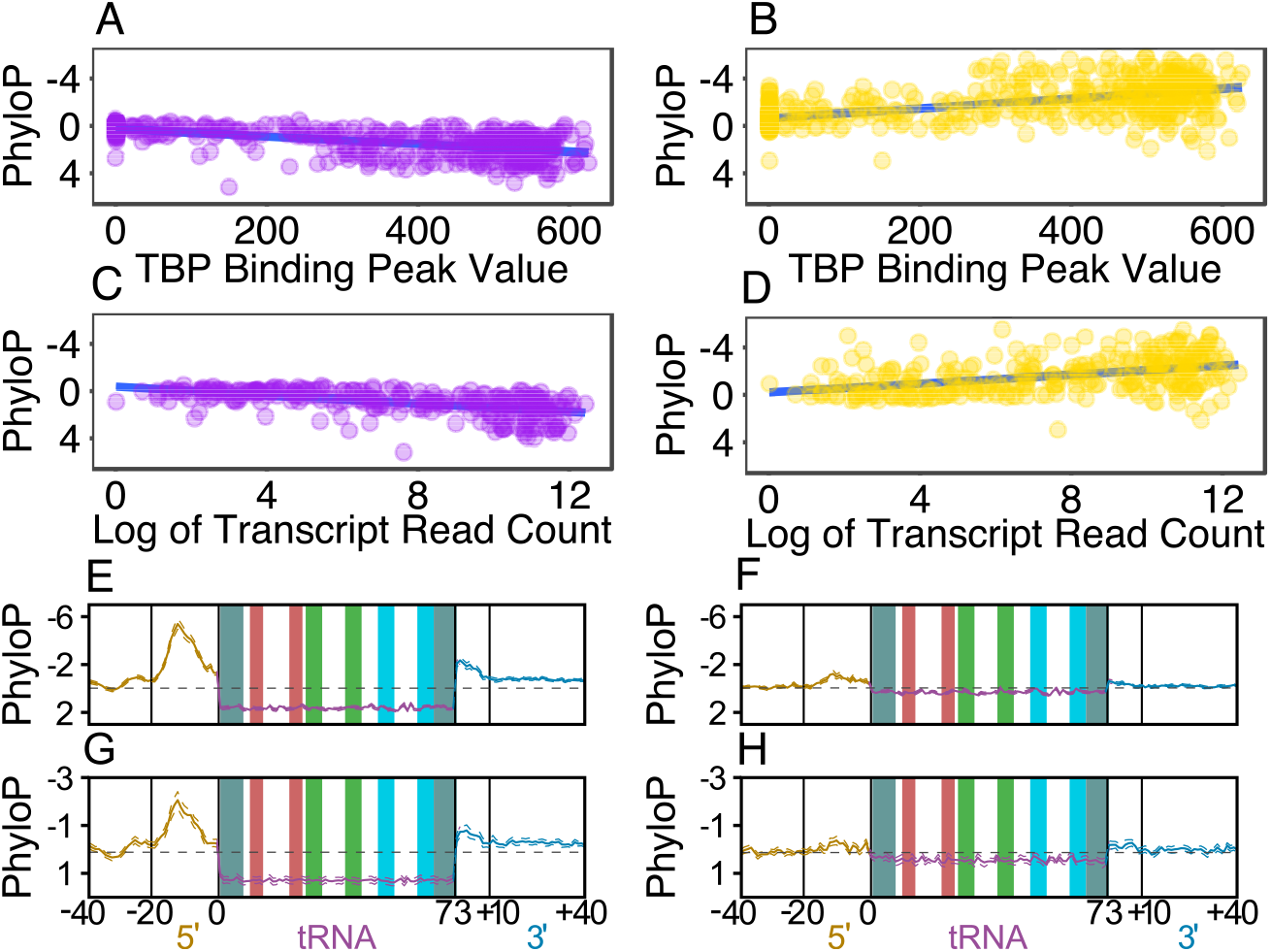
tRNA expression is significantly correlated to both tRNA conservation and flanking region divergence. TATA-Binding Protein (TBP) peak value (expression) plotted versus PhyloP score (conservation) for each (**A**) mature tRNA and (**B**) adjacent inner 5’ flanking region. Log of the HEK293T cell DM-tRNA-seq read count (expression; 32) plotted versus PhyloP score (conservation) for each (**C**) gene encoding a *unique* mature tRNA sequence and (**D**) corresponding inner 5’ flanking region. Both TBP occupancy and transcript abundance are greater for highly conserved mature tRNA loci (**A,C**) and those with the most divergent flanks (**B,D**). Plotted as in Fig. 1, human tRNA loci that are separated into (**E**) active versus (**F**) inactive groups show characteristic differences seen in (**A-D**). Mouse tRNA loci split into (**G**) active versus (**H**) inactive groups show a strikingly similar pattern as seen in human (**A-F**).

We find that active tRNA genes are significantly more conserved than inactive tRNA loci (Mann-Whitney U, p < 8.40e-53), and the flanking regions of active tRNAs are significantly more *divergent* than the flanking regions of inactive tRNAs (p < 7.98e-61). The peak measure of divergence between human and *M. mulatta* tRNA genes in the inner 5’ flanking regions is roughly five times greater in active tRNAs than in inactive tRNAs (Fig. 2E,F). Active tRNAs also have significantly more low-frequency SNPs per site in human populations than inactive tRNAs across the entire locus, including the tRNA and flanking regions (p < 3.72e-36; SI Appendix, Fig. S3). Inactive tRNAs are still significantly more conserved (p < 2.02e-12) and polymorphic (p < 0.007) than the untranscribed reference regions, and their flanks are significantly more divergent than the reference regions (p < 1.36e-16).

That the peak in both divergence and polymorphism in all species is consistently 12 to 15 nucleotides upstream of the mature tRNA sequence is curious. At the most divergent position, 55% of all tRNA loci differ between human and *M. mulatta* and 15% of human tRNA loci have a low-frequency SNP (SI Appendix, Fig. 3A). Furthermore, virtually all active tRNA loci differ at this nucleotide between human and *M. mulatta,* and 25% have a low-frequency SNP at this site (SI Appendix, Fig. S3B). This implies that this region either does not face uniform selective pressures or is not uniformly vulnerable to TAM. While distant flanking sequences can affect tRNA expression in yeast (34), few studies have shown that flanking regions affect expression in higher eukaryotes (35). Transcription initiation is long relative to elongation (36, 37), which may lead to prolonged isolation of the non-template DNA strand at the initiation site and increased vulnerability to TAM. A poised initiation complex might also increase the likelihood of collisions between Pol3 and the replication fork (14). Thus, frequent initiation at highly transcribed tRNA loci may contribute to the non-uniform pattern of variation.

**Fig. 3.**
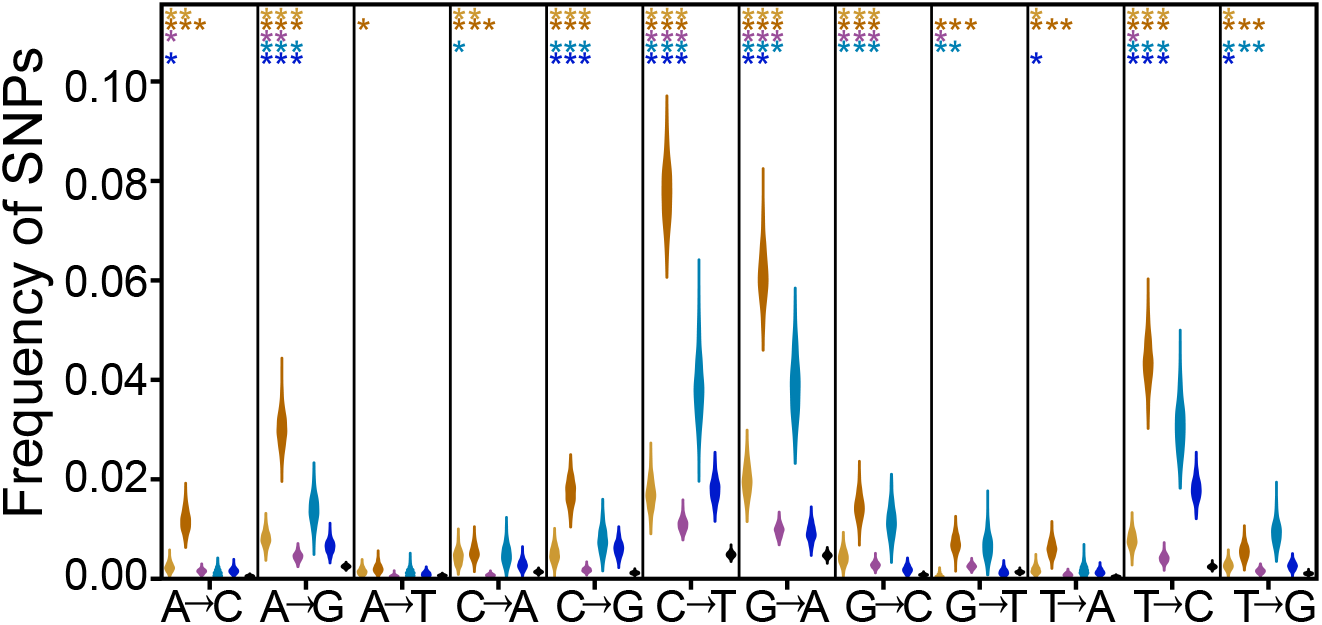
SNP classes most common in regions affected by TAM are also most common at tRNA loci. Distribution of each class of low-frequency polymorphisms by region across all human tRNAs. Significance levels of Fisher’s exact tests comparing the SNP distribution within each region of the tRNA and flank (outer 5’ flank in yellow, inner 5’ flank in orange, tRNA in purple, inner 3’ flank in cyan, outer 3’ flank in blue) to that of the untranscribed reference region (black) represented by stars. One star represents a p value ≤ 0.05, two stars ≤ 0.005, and three stars ≤ 0.0005.

This may also explain the increased variation in the outer 3’ flank relative to the outer 5’ flank, as positioning of downstream transcription termination sites is varies among tRNA genes (2, 38), whereas transcription start site positions are more consistent. While most tRNAs do not have clear TATA boxes, the TATA-Binding Protein (TBP) still binds to the DNA duplex approximately 25 nucleotides upstream of the tRNA (39), which coincides with a decrease in variability. Furthermore, while both flanking regions for many other Pol3-transcribed genes are divergent, the 5’ flanking regions are generally more divergent than the 3’ flanking regions, suggesting that the underlying mechanism is not tRNA-specific (SI Appendix, Dataset S1). However, additional studies are necessary to support the assertion that this pattern is due to transcription.

Two orthogonal analyses strengthen the observed correlations between gene expression and variation at tRNA loci. First, we find a significant correlation between the TBP intensity peaks (40–42) and conservation of the mature tRNA sequence (Spearman’s rho = 0.64, p < 2.2e-16) across all human tRNAs and the opposite relationship in the flanking regions (Spearman’s rho = -0.64, p < 2.2e-16; Fig. 2). TBP ChIP-Seq data directly reflects transcriptional activity for each locus, as its occupancy is significantly correlated with and required for transcription (20, 43–47). Second, mature tRNA sequence read counts are strongly correlated with tRNA conservation (Spearman’s rho = 0.18, p < 0.001) and flanking region divergence (Spearman’s rho = -0.61, p < 2.2e-16; Fig. 2; SI Appendix, Fig. S4). These read counts were collected from a single human embryonic kidney cell line by Zheng et al. (32) using DM-tRNA-seq, a specialized tRNA sequencing method. These correlations are consistent with the idea that more highly transcribed tRNAs vary more in their flanking regions.

### Patterns of divergence and conservation can be leveraged to predict tRNA gene expression

Regardless of how tRNA expression is measured, we find highly significant correlations between gene expression and tRNA sequence conservation. The consistency of these correlations across methods of measurement and across species indicates that it may be possible to predict relative tRNA, with DNA sequence conservation patterns and other correlates of tRNA transcriptional activity (e.g., tRNAscan-SE bit scores). Indeed, active and inactive tRNAs are largely distinguishable using only flank and gene PhyloP data (SI Appendix, Fig. S5). As sequencing technology becomes more accessible, predicting tRNA gene expression levels through analysis of DNA data is enticing. Such a model could make future tRNA gene annotation more detailed and cost-effective.

### Variation patterns observed at tRNAs are not observed in most other gene families

Applicability of this proposed tool is likely best suited for tRNAs, other Pol3 genes, and unique classes of highly expressed protein coding genes such as histones. Among the histone protein coding genes less than 1,000 nucleotides in length, the average PhyloP score per nucleotide across the coding sequence and flanking regions are 3.449 and -2.052, respectively, comparable to tRNA loci (SI Appendix, Fig. S6). In contrast, most genes transcribed by RNA Pol2 do not appear to be good targets (SI Appendix, Dataset S1). For example, ribosomal proteins are very highly transcribed (48) and have well conserved exons, but their introns and flanking regions are not as divergent as tRNA flanking regions (28, 49). tRNAs are likely ideal for studying TAM because they have predictable transcript start and end sites, internal promoters, and high transcription rates.

### Patterns of low-frequency SNPs are consistent with transcription-associated mutagenesis (TAM)

In TAM, repair pathways activated in response to deaminations lead to excess conversions between guanine and adenine and between thymine and cytosine on the coding strand (9, 17). Across all tRNA loci, we found that the most common low-frequency SNPs are C→T and G→A, and that these mutations are significantly more common in both tRNA genes and flanking regions, relative to untranscribed reference regions (Fisher’s exact test, p < 0.05 for all comparisons; Fig. 3). Removal of CpG sites (50) does not significantly affect these results. The relative excesses of these SNPs are much more pronounced in active tRNA loci than in inactive tRNA loci (SI Appendix, Fig. S7A,B). These results suggest that deamination of the non-coding strand due to TAM and the DNA repair mechanisms acting in response to deamination are especially common at these loci (9, 17, 19).

It is difficult to discern whether this increased prevalence is due to TAM or selection to preserve structural integrity of the tRNA. To preserve tRNA secondary structure, we expect transition mutations (e.g., A-U to G-U base pairs, C-G to U-G base pairs) to be more common than transversions, as they should disrupt stem helices less often. However, the mutational skew expected of regions affected by TAM is stronger in regions flanking tRNAs. Transcription initiation is relatively long compared to elongation (36, 37), which might contribute to increased mutagenesis by APOBEC enzymes, more collisions (14) or double-stranded breaks. However, divergence at tRNA flanking regions is correlated with divergence at introns in both human (SI Appendix, Fig. S1B, Spearman’s rank, rho = 0.734, p < 5.58e-6) and mouse (SI Appendix, Fig. S1D, rho = 0.733, p < 5.24e-4), indicating similar mutation rates across tRNA loci. Our results therefore suggest that TAM drives the excess of transitions among low-frequency SNPs across tRNA loci.

### tRNA flanking region variation in other model organisms is consistent with variation observed in humans

To confirm that our results are not restricted to humans, we also analyzed tRNAs in *Mus musculus, Drosophila melanogaster* and *Arabidopsis thaliana.* We find similar patterns of sequence conservation of tRNA loci in each, when measuring PhyloP or divergence to outgroups (SI Appendix, Fig. S8). The 5’ flanks are consistently more divergent than the 3’ flanks, and the most divergent sites are roughly 10-15 bases upstream of the tRNA in all species. We also used chromatin-IP data across nine mouse tissues to classify mouse tRNAs based on their expression (51). Active mouse tRNAs are more strongly conserved than their inactive counterparts (Mann-Whitney U, p < 1.81e-19), and their flanks are more divergent (p < 7.04e-22, Fig. 2G,H), consistent with our results from the human data (Fig. 2E,F). Active mouse tRNAs also have more low-frequency SNPs in their flanking regions than inactive mouse tRNAs (p < 2.23e-4; SI Appendix, Fig. S9). Such consistency suggests that a shared underlying molecular mechanism drives these patterns of sequence variation.

Low-frequency SNPs in the tRNA gene sequences also follow similar qualitative patterns to the human data. We observe excess transitions in all species studied (SI Appendix, Fig. S10), and active mouse tRNAs show a greater excess of low-frequency transitions than do inactive mouse tRNAs (SI Appendix, Fig. S7C,D). However, these patterns vary across species (Fig. 4B-D). For example, in mouse, tRNA genes have more low-frequency SNPs than the untranscribed reference regions (Fig. 4B), but the opposite is true in *D. melanogaster* (Fig. 4D). Low-frequency SNPs are thought not to be strongly affected by selection (29), but selection is more efficient in species with greater effective populations sizes (Fig. 4A). Effective population size (52–55) and tRNA copy number vary across species, and because the sample sizes and data quality differ between population samples, these differences may be attributable to differences in the impact of selection or in ascertainment of low-frequency variation.

**Fig. 4.**
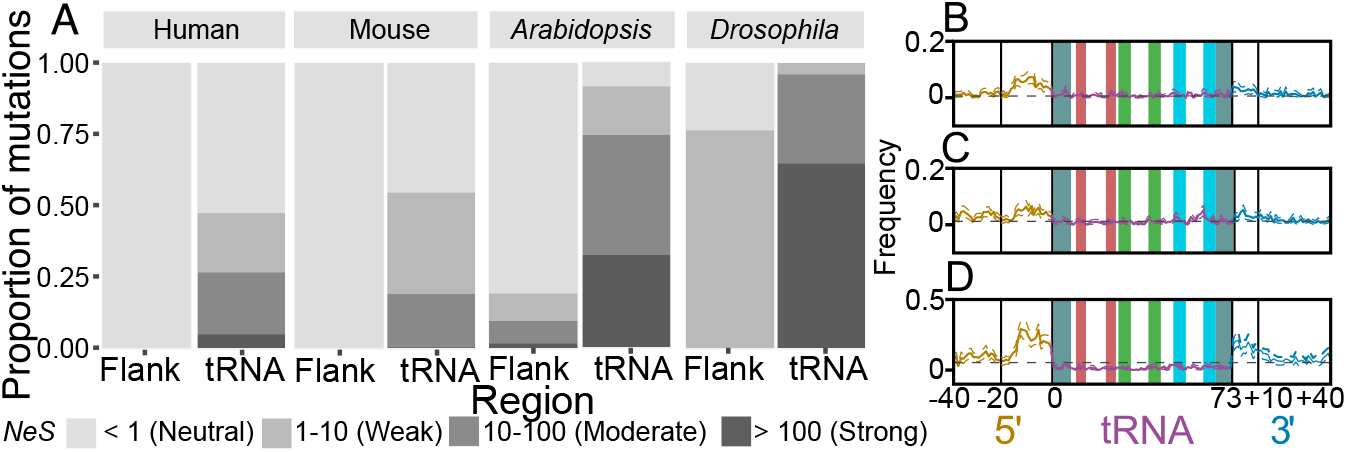
Estimated Distribution of Fitness Effects (DFE) indicates that high proportions of deleterious mutations in tRNAs are under strong selection. (**A**) Estimated DFE of new deleterious mutations for tRNA genes and inner 3’ flanking regions shown in human, mouse, *A. thaliana* and *D. melanogaster.* Proportions of deleterious mutations shown for each bin of purifying selection strength, estimated on a scale of N_e_S, where N_e_ is the effective population size and S is the strength of selection. Species are arranged by increasing N_e_. Low-frequency SNPs plotted as in Fig. 1C for (**B**) mouse, (**C**) *A. thaliana*, and (**D**) *D. melanogaster.*

### Functional tRNA sequences experience strong purifying selection in all species studied

Our analysis of the distribution of fitness effects (DFE) of deleterious mutations demonstrates that tRNAs evolve under strong purifying selection in all analyzed species. In contrast, regions flanking tRNAs are inferred to be either neutral or subject to weak selection (N_e_S < 10; Fig. 4A). Our estimates of the proportions of new mutations falling into each N_e_S range of the DFE for tRNAs indicate far fewer nearly neutral mutations (N_e_S < 1) and substantially more strongly deleterious mutations (N_e_S > 100 in *D. melanogaster* and *A. thaliana* than in the human or mouse populations (Fig. 4A). Given that estimates of N_e_ in humans (7,000; 52) and mouse (25,000-120,000; 53) are substantially lower than in *A. thaliana* (300,000; 54) and *D. melanogaster* (> 1,000,000; 55), this difference in strength of selection may partially reflect differences in N_e_, and might explain the differences in low-frequency SNPs in tRNA loci across species (Fig. 4B-D). In turn, this might indicate that the strength of purifying selection, independent of effective population size, at tRNA loci is consistent across diverse species.

The strength of selection across species may also reflect the number of unique tRNA gene sequences in each genome. For example, roughly half of all human tRNA genes have unique sequences, but the majority of *D. melanogaster* tRNAs have identical copies (2). tRNAs with the same anticodon but different sequences may have different functions, and that this may affect strength of selection at each locus as well. Indeed, a significantly greater proportion of sites are invariant (Fisher’s exact test, p < 7.50e-5) and fewer sites are divergent (p < 3.85e-8) in active single-copy human tRNA genes than in active multi-copy human tRNA genes. We observe the same patterns in the inner 5’ (p < 5.87e-5; p < 0.025) and inner 3’ (p < 8.90e-5; p < 4.04e-4) flanks of active tRNA genes, suggesting increased transcription of active multi-copy tRNA genes. However, little SNP data is available for multi-copy tRNAs compared to single-copy tRNAs, limiting our ability to identify consistent differences among tRNA subgroups.

### tRNA loci contribute disproportionately to mutational load

Our discovery of a highly elevated mutation rate at tRNA loci suggests that tRNA genes may contribute disproportionately to mutational load, the reduction in individual fitness due to deleterious mutations (56, 57). To estimate the relative mutation rates at active tRNA loci, we calculated the average ratios of θ for the inner 3’ and 5’ flanking regions of active human tRNA genes to the untranscribed reference regions using the approach of Messer (29; see Methods). We estimate θ in the flanking regions instead of the tRNAs because strong selection can cause underestimation of θ (29), and our results indicate that active human tRNAs are subject to strong selection while the flanking regions are likely selectively neutral (Fig. 4A). We therefore estimate that the mutation rate is between 7.24 (inner 3’; 95% confidence interval 7.12-7.33) and 10.36 (inner 5’; 10.16-10.41) times greater at tRNA loci than the genome-wide average. Given that there are 25,852 base pairs of active human tRNA sequence, and using 1.45e-8 as the genome-wide mutation rate (58), we estimate that U (the genome-wide rate of deleterious mutation per diploid genome) contributed by tRNAs is between 0.0054 and 0.0078. Since active tRNAs make up only 0.0009% of the human genome (2), this implies that mutations in tRNAs contribute disproportionately to mutational load. Our findings highlight that mutations at tRNA loci are likely an important source of fitness and disease variation in human populations.

## Conclusions

Our findings demonstrate the exceptional transcription rates of tRNA genes causes a similarly substantial increase in mutation rates through TAM. Our results are consistent across a broad range of taxonomically diverse species, indicating that elevated mutation rates due to TAM and strong purifying selection are widespread, and may be a good predictor of relative tRNA gene transcription levels. The conflict between extreme TAM and consequent strong purifying selection at tRNA loci is potentially an unappreciated source of genetic disease, and may have a profound impact on human fitness, yet to be fully addressed.

## Materials and Methods

### Defining tRNA loci and flanking regions

We used tRNA coordinates from GtRNAdb (2) for the human, *M. musculus, D. melanogaster,* and *A. thaliana* genomes. For each species, we defined untranscribed reference regions by searching 10 kilobases upstream of each tRNA and selecting a 200-nucleotide tract. If this tract was within a highly transcribed region of the genome (based on genome-wide chromatin-IP data; 33), overlapped a conserved element (defined as a region with a phastCons log odds score greater than 0; 28), was within 1,000 nucleotides of a known gene (49), or overlapped a reference region assigned to another tRNA, we selected a new tract 1,000 bases further upstream, and repeated until we found an acceptable region. For the mouse genome, we checked known genes, previously assigned reference regions, and conserved elements. For the *D. melanogaster* and *A. thaliana* genomes, we began our searches only 1,000 bases upstream of each tRNA, and searched for 200-nucleotide tracts that were at least 100 nucleotides away from any annotated genetic element (59, 60) due to the high functional densities of these species’ genomes.

For each tRNA in all species, we defined the inner 5’ flank as the 20 bases immediately upstream of the 5’ end of the tRNA gene on the coding strand, and the outer 5’ flank as the 20 bases directly upstream of the inner 5’ flank. The inner 3’ flank refers to the 10 bases downstream of the tRNA gene, and the outer 3’ flank refers to the 30 bases downstream of these 10 bases. We made these decisions based on inflection points in our data, as the flanking regions up to 20 bases upstream and 10 bases downstream of tRNA genes have less variation. Transcription usually ends about 10 bases downstream of tRNA genes (38).

### Classifying tRNAs based on breadth of expression

The Roadmap Epigenomics Consortium compiled genome-wide epigenomic data across 127 human tissues and cell lines in order to characterize the chromatin state across the genome (33). We analyzed the regions surrounding each tRNA in each epigenome sample, and used clustering to classify each genomic region according to its most common epigenomic state. We classified all human tRNAs based on the epigenomic state annotation in the genome. In the corresponding model, regions in state 1 are likely to be transcribed. The 342 tRNAs in state 1 in at least four of the 127 tissues analyzed are “active tRNAs”, and we consider the remaining 254 tRNAs “inactive”. To classify mouse tRNAs, we used a 15-state Hidden Markov Model based on chromatin-IP data in which states 5 and 7 corresponded to regions near active promoters (51). We considered the 272 tRNAs in genomic regions annotated as state 5 or 7 in at least 3% of tissues “active” and the remaining 188 tRNAs “inactive”.

### Aligning tRNAs

We aligned all tRNAs across all species using covariance models (61) and assigned coordinates to each position in each tRNA and flank based on the Sprinzl numbering system (31). We averaged the PhyloP, divergence and low-frequency SNP data for all sites assigned to the same Sprinzl coordinate for their respective tRNA loci. Because some tRNAs have variations in structure (2), this alignment was necessary for position-wise comparisons between tRNAs. We filtered tRNAs with fewer than 50 aligned bases from our analyses. If a conserved element (regions with a phastCons log odds score greater than 0; 28) was present 4-10 bases up or downstream of a tRNA, the tRNA was excluded from our analyses, as these regions might contribute to the secondary structure of mature tRNAs and be subject to anomalous levels of selection. We also excluded nuclear-encoded mitochondrial tRNA genes.

### Parsing variation data

We analyzed human variation data from the African superpopulation of 661 humans from Phase 3 of the 1000 Genomes Project (62). We acquired *D. melanogaster* variation data for the Siavonga, Zambia populations from the Drosophila Genome Nexus Database (59, 60). We obtained *M. musculus* and *A. thaliana* data from Waterston, et al. (63) and the Arabidopsis Genome Initiative (64), respectively. All nonhuman data were aligned and genotypes curated as described in Corbett-Detig, et al. (65).

Within each gene, flank or reference region, we considered positions with minor allele frequencies between 0 and 0.05 to be low-frequency SNPs. We also determined the frequency each class of mutations (e.g. A→G) within each region of each tRNA locus where the identity of each base is defined according to the coding strand sequence. We found the frequency of divergences and low-frequency SNPs by position across all tRNAs and flanking regions. For conservation studies across multiple species, we used the PhyloP track (28) from the UCSC Genome Browser (49) and calculated the average score for each position within the tRNAs and flanking regions. No PhyloP data was available for *A. thaliana* (28). For direct comparisons between the species of interest and an outgroup, we used the Multiz track from the UCSC Table Browser (66) and the Stitch MAFs tool from Galaxy (67) to create sequence alignments. Details are available in the SI Appendix.

### Transcription factor binding

The ENCODE Project Consortium used ChIP-Seq data to identify binding regions for regulatory factors (40–42), including the TATA-binding protein (TBP) and Pol3 transcription factors in the human genome (20). These data were taken from the UCSC Genome Browser (49). The intensity of a given peak correlates with a greater frequency of transcription factor binding to that region. For each human tRNA, we found the strongest TBP peak in the 50 base pairs immediately upstream of the tRNA, across the GM12878, H1-hESC, HeLa-S3, HepG2 and K562 cell lines. We also calculated the average PhyloP score across the flanking regions for each tRNA (28), and used Spearman’s rank correlation test on these data.

### Correlating variation to cell-line read counts

Zheng, et al. (32) used demethylation sequencing to detect tRNAs within human embryonic kidney (HEK293T) cells (32, 68). We used Spearman’s rank correlation tests to correlate mature tRNA transcript read counts and tRNA and flanking region conservation. Because Zheng, et al. (32) sequenced mature tRNAs, which are often encoded by multiple genes, we excluded identical genes to control for the correlation between gene copy-number and overall expression (Fig. 2C,D; 32, 34). Separately, we summed the average PhyloP scores at these loci and correlating to total tRNA read counts (SI Appendix, Fig. S4).

### Estimating the distribution of fitness effects

We estimated the distribution of fitness effects (DFE) for each species using the method of Keightley et al. (69) and the DFE-α software. See SI Appendix for details.

### Estimating mutation rate in active tRNA genes

We used the equation θ(k) = kG^k^ defined in Messer 2009 (29) to estimate the mutation rate at active tRNA loci. We calculated the ratios of θ in active tRNA flanking regions to θ in the reference regions for k = 1,2,3 and bootstrapped by tRNA loci to calculate 95% confidence intervals. See SI Appendix for more details.

## Acknowledgments

We thank Craig Mello for helpful discussions and input on this project, Brian Lin for the Fig. 1 legend, and Andrew Holmes for assisting with mouse data analysis. We thank the Corbett and Lowe Labs for suggestions and feedback. This work was supported by a grant from the National Human Genome Research Institute, National Institutes of Health (2R01HG006753-04A1 to T.L.). B.T. was funded by an NIH training grant (T32 HG008345).

